# Deep Learning Interpretation of Echocardiograms

**DOI:** 10.1101/681676

**Authors:** Amirata Ghorbani, David Ouyang, Abubakar Abid, Bryan He, Jonathan H. Chen, Robert A. Harrington, David H. Liang, Euan A. Ashley, James Y. Zou

## Abstract

Echocardiography uses ultrasound technology to capture high temporal and spatial resolution images of the heart and surrounding structures and is the most common imaging modality in cardiovascular medicine. Using convolutional neural networks on a large new dataset, we show that deep learning applied to echocardiography can identify local cardiac structures, estimate cardiac function, and predict systemic phenotypes that modify cardiovascular risk but not readily identifiable to human interpretation. Our deep learning model, EchoNet, accurately identified the presence of pacemaker leads (AUC = 0.89), enlarged left atrium (AUC = 0.85), normal left ventricular wall thickness (AUC = 0.75), left ventricular end systolic and diastolic volumes(*R*^2^ = 0.73 and *R*^2^ = 0.68), and ejection fraction (*R*^2^ = 0.48) as well as predicted systemic phenotypes of age (*R*^2^ = 0.46), sex (AUC = 0.88), weight (*R*^2^ = 0.56), and height (*R*^2^ = 0.33). Interpretation analysis validates that EchoNet shows appropriate attention to key cardiac structures when performing human-explainable tasks and highlight hypothesis-generating regions of interest when predicting systemic phenotypes difficult for human interpretation. Machine learning on echocardiography images can streamline repetitive tasks in the clinical workflow, standardize interpretation in areas with insufficient qualified cardiologists, and more consistently produce echocardiographic measurements.

## Introduction

Cardiovascular disease has a substantial impact on overall health, well-being, and life-expectancy. In addition to being the leading cause of mortality for both men and women, cardiovascular disease is responsible for 17% of the United States’ national health expenditures.^1^ Even as the burden of cardiovascular disease is expected to rise with an aging population^1^, there continues to be significant racial, socioeconomic, and geographic disparities in both access to care and disease outcomes.^2, 3^ Variation in access to and quality of cardiovascular imaging has been linked to disparities in outcomes.^3, 4^ It has been hypothesized that automated image interpretation can enable more available and accurate cardiovascular care and begin to alleviate some of the disparities in cardiovascular care.^5, 6^ The application of machine learning in cardiology is still in its infancy, however there is significant interest in bringing neural network based approaches to cardiovascular imaging.

Machine learning has transformed many fields, ranging from image processing and voice recognition systems to super-human performance in complex strategy games.^7^ Many of the biggest recent advances in machine learning come from computer vision algorithms and processing image data with deep learning.^8–11^ Recent advances in machine learning suggest deep learning can identify human-identifiable characteristics as well as phenotypes unrecognized by human experts.^12, 13^ Efforts to apply machine learning to other modalities of medical imaging have shown promise in computer-assisted diagnosis.^12–16^ Seemingly unrelated imaging of individual organ systems such as fundoscopic retina images can predict systemic phenotypes and predict cardiovascular risk factors.^12^ Additionally, deep learning algorithms perform well in risk stratification and classification of disease.^14, 16^. Multiple recent medical examples outside of cardiology show convolutional neural network algorithms can match or even exceed human experts in identifying and classifying diseases.^13, 14^

Echocardiography is a uniquely well-suited approach for the application of deep learning in cardiology. The most readily available and widely used imaging technique to assess cardiac function and structure, echocardiography combines rapid image acquisition with the lack of ionizing radiation to serve as the backbone of cardiovascular imaging.^4, 17^ Echocardiography is both frequently used as a screening modality for healthy, asymptomatic patients as well as in order to diagnose and manage patients with complex cardiovascular disease.^17^ For indications ranging from cardiomyopathies to valvular heart diseases, echocardiography is both necessary and sufficient to diagnose many cardiovascular diseases. Despite its importance in clinical phenotyping, there is variance in the human interpretation of echocardiogram images that could impact clinical care.^18–20^ Formalized training guidelines for cardiologists recognize the value of experience in interpreting echocardiogram images and basic cardiology training might be insufficient to interpret echocardiograms at the highest level.^21^

Given the importance of imaging to cardiovascular care, an automated pipeline for standardizing and interpreting cardiovascular imaging can improve peri-operative risk stratification, manage the cardiovascular risk of patients with oncologic disease undergoing chemotherapy, and aid in the diagnosis of cardiovascular disease.^1, 22, 23^ While other works applying machine learning to medical imaging required re-annotation of images by human experts, the clinical workflow for echocardiography inherently includes many measurements and calculations and often is reported through structured reporting systems. The ability to use previous annotations and interpretations from clinical reports can greatly accelerate adoption of machine learning in medical imaging. Given the availability of previously annotated clinical reports, the density of information in image and video datasets, and many available machine learning architectures already applied to image datasets, echocardiography is a high impact and highly tractable application of machine learning in medical imaging.

### Related works

Current literature have already shown that it is possible to identify standard echocardiogram views from unlabeled datasets.^5, 6, 24^ Previous works have used convolutional neural networks (CNNs) trained on images and videos from echocardiography to perform segmentation to identify cardiac structures and derive cardiac function. In this study, we extend previous analyses to show that EchoNet, our deep learning model using echocardiography images, local cardiac structures and anatomy, estimate volumetric measurements and metrics of cardiac function, and predict systemic human phenotypes that modify cardiovascular risk. Additionally, we show the first application of interpretation frameworks to understand deep learning models from echocardiogram images. Human identifiable features, such as the presence of pacemaker and defibrillator leads, left ventricular hypertrophy, and abnormal left atrial chamber size identified by our convolutional neural network were validated using interpretation frameworks to highlight the most relevant regions of interest. To the best of our knowledge, we develop the first deep learning model that can directly predict age, sex, weight and height from echocardiogram images and use interpretation methods to understand how the model predicts these systemic phenotypes difficult for human interpreters.

## Results

**Figure 1.**
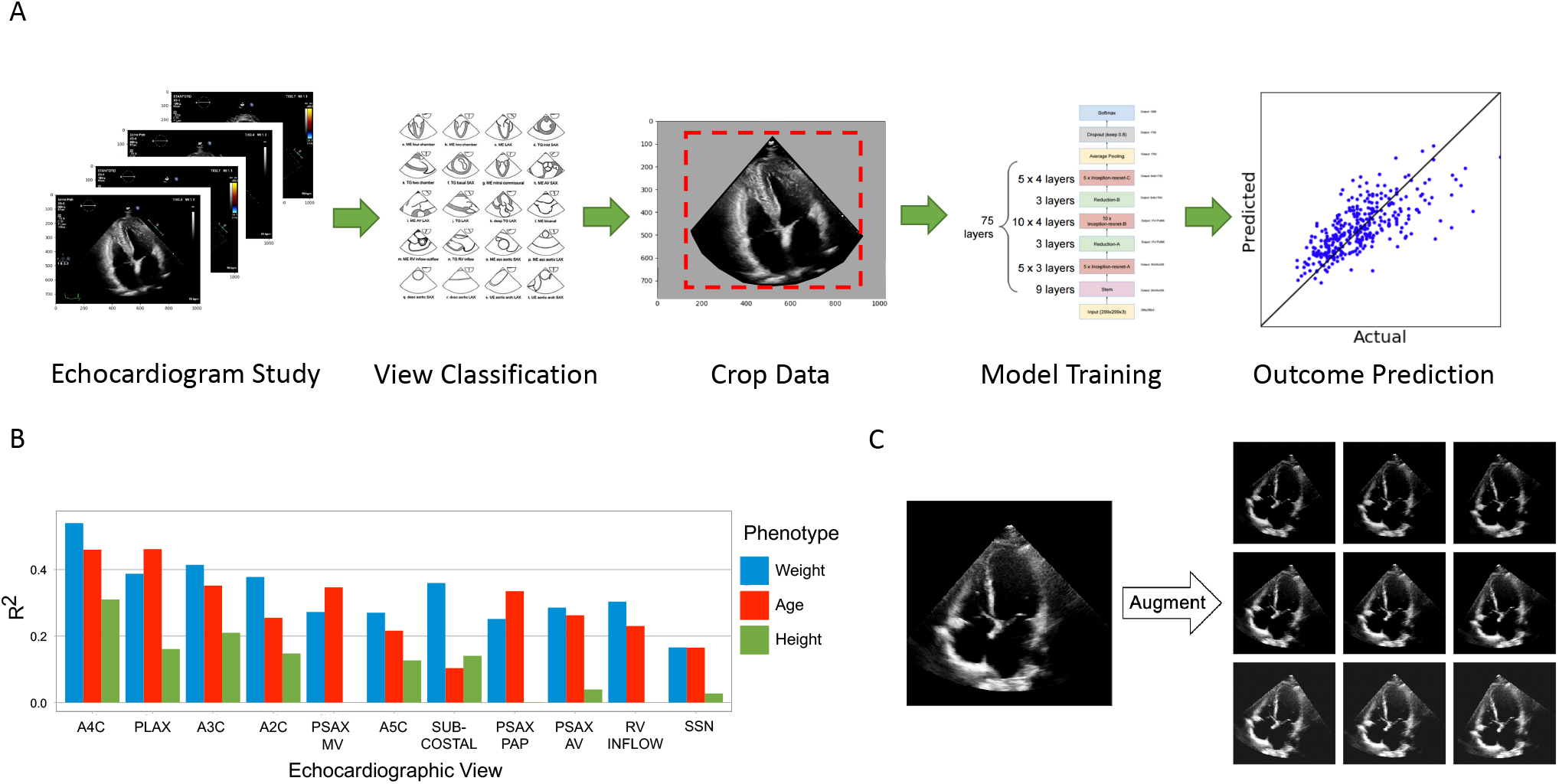
A. EchoNet workflow for image selection, cleaning, and model training. B. Comparison of model performance with different cardiac views as input. C. Examples of data augmentation. The original frame is rotated (left to right) and its intensity is increase (top to bottom) as augmentations.

We trained a convolutional neural network model on a data set of more than 2.6 million echocardiogram images from 2,850 patients to identify local cardiac structures, estimate cardiac function, and predict systemic risk factors (Fig. **??**). Echocardiogram images, reports, and measurements were obtained from an accredited echocardiography lab of a large academic medical center (Table 1). Echocardiography visualizes cardiac structures from various different orientations and geometries, so images were classified by cardiac view to homogenize the input data set. Echocardiogram images were sampled from echocardiogram videos, pre-processed by de-identifying the images, and cropped to eliminate information outside of the scanning sector. These processed images were used to train EchoNet on the relevant medical classification or prediction task.

**Table 1.** Baseline characteristics of patients in the development and validation datasets

| Characteristics | Complete Data |  | A4C View Data |  |
| --- | --- | --- | --- | --- |
|  | Train Data | Test Data | Train Data | Test Data |
| Number of Patients | 2850 | 373 | 2546 | 337 |
| Number of Images | 1,624,780 | 169,880 | 172,080 | 21,540 |
| Sex (% Male) | 52.4% | 52.8% | 52.2% | 53.7% |
| Age: mean, years (std) | 61.3 (17.2) | 62.8 (16.8) | 61.1 (17.1) | 63.2 (16.9) |
| Weight: mean, Kg (std) | 78.8 (22.7) | 78.9 (20.8) | 78.0 (21.7) | 78.5 (20.2) |
| Height: mean, m (std) | 1.69 (0.11) | 1.69 (0.11) | 1.69 (0.12) | 1.69 (0.11) |
| BMI: mean (std) | 27.3 (6.7) | 27.5 (6.5) | 27.1 (6.5) | 27.3 (6.1) |
| Pacemaker or Defibrillator Lead (% Present) | 13.2 | 14.7 | 13.1 | 15.1 |
| Severe Left Atrial Enlargement (% Present) | 17.2 | 20.3 | 18.0 | 21.9 |
| Left ventricular hypertrophy (% Present) | 33.3 | 38.0 | 32.7 | 37.9 |
| End Diastolic Volume, mL: mean (std) | 94.3 (47.2) | 94.6 (13.0) | 95.1 (48.2) | 96.9 (48.0) |
| End Systolic Volume, mL: mean (std) | 45.6 (38.3) | 46.2 (36.1) | 46.0 (39.3) | 47.0 (36.6) |
| Ejection Fraction: mean (std) | 55.2 (12.3) | 54.7 (13.0) | 55.1 (12.2) | 54.8 (13.1) |

### Predicting anatomic structures and local features

A standard part of the clinical workflow of echocardiography interpretation is the identification of local cardiac structures and characterization of its location, size, and shape. Local cardiac structures can have significant variation in image characteristics, ranging from bright echos of metallic intracardiac structures to dark regions denoting blood pools in cardiac chambers. As our first task, we trained EchoNet on three classification tasks frequently evaluated by cardiologists that rely on recognition of local features (Fig. 2). Labels of the presence of intracardiac devices (such as catheters, pacemaker, and defibrillator leads), severe left atrial dilation, and normal left ventricular wall thickness were extracted from the physician-interpreted report and used to train EchoNet on unlabeled apical-4-chamber input images. The presence of a pacemaker lead was predicted with high accuracy (AUC of 0.89, F1 score of 0.73), followed by the identification of a severely dilated left atrium (AUC of 0.85, F1 score of 0.68), and normal left ventricular wall thickness (AUC of 0.75, F1 score of 0.56). To understand the model’s predictions, we used gradient-based sensitivity map methods^25^ to identify the regions of interest for the interpretation and show that EchoNet highlights relevant areas that correspond to intracardiac devices, the left atrium, and the left ventricle respectively. Models’ prediction robustness was additionally examined with direct input image manipulations, including occlusion of human recognizable features, to validate that EchoNet arrives at its predictions by focusing on biologically plausible regions of interest.^26^ For example, in the frames in Figure 2 with pacemaker lead, when we manually mask the lead in the frame, EchoNet changs its prediction to no pacemaker.

**Figure 2.**
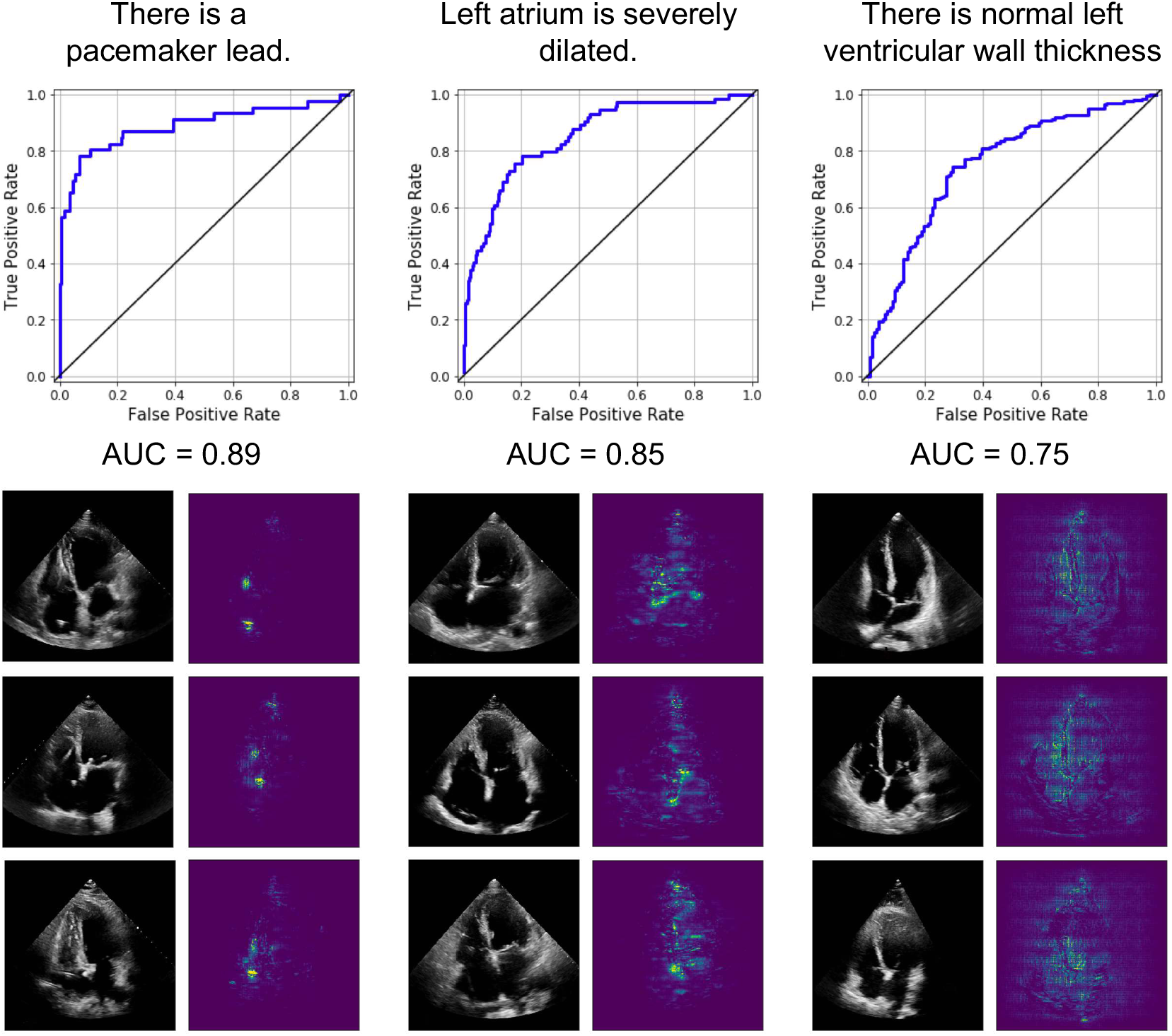
EchoNet performance and interpretation for three clinical interpretations of local structures and features. For each task, representative positive examples are shown side-by-side with regions of interest from the respective model.

### Predicting cardiac function

Quantification of cardiac function is a crucial assessment addressed by echocardiography. However it has significant variation in human interpretation^18, 19^. The ejection fraction, a measure of the volume change in the left ventricle with each heart beat, is a key metric of cardiac function, but its measurement relies on the time-consuming manual tracing of left ventricular areas and volumes at different times during the cardiac cycle. We trained EchoNet to predict left ventricular end systolic volume (ESV), end diastolic volume (EDV), and ejection fraction from sampled apical-4-chamber view images (Fig. 3). Left ventricular ESV and EDV were accurately predicted. For the prediction of ESV, an *R*^2^ score of 0.74 and mean absolute error (MAE) of 13.3 mL was achieved versus MAE of 25.4 mL if we use mean prediction which is to predict every patient’s ESV as the average ESCV value of patients. The result for the EDV prediction was an *R*^2^ score of 0.70 and MAE of 20.5 mL (mean prediction MAE = 35.4 mL). Conventionally, ejection fraction is calculated from a ratio of these two volumetric measurements, however calculated ejection fraction from the predicted volumes were less accurate (Fig. 3C) than EchoNet trained directly on the ejection fraction (Fig. 3D). Using the trained EchoNet, an *R*^2^ score of 0.50 and MAE of 7.0% is achieved (MAE of mean prediction = 9.9%). For each model, interpretation methods show appropriate attention over left ventricle as the region of interest to generate the predictions.

**Figure 3.**
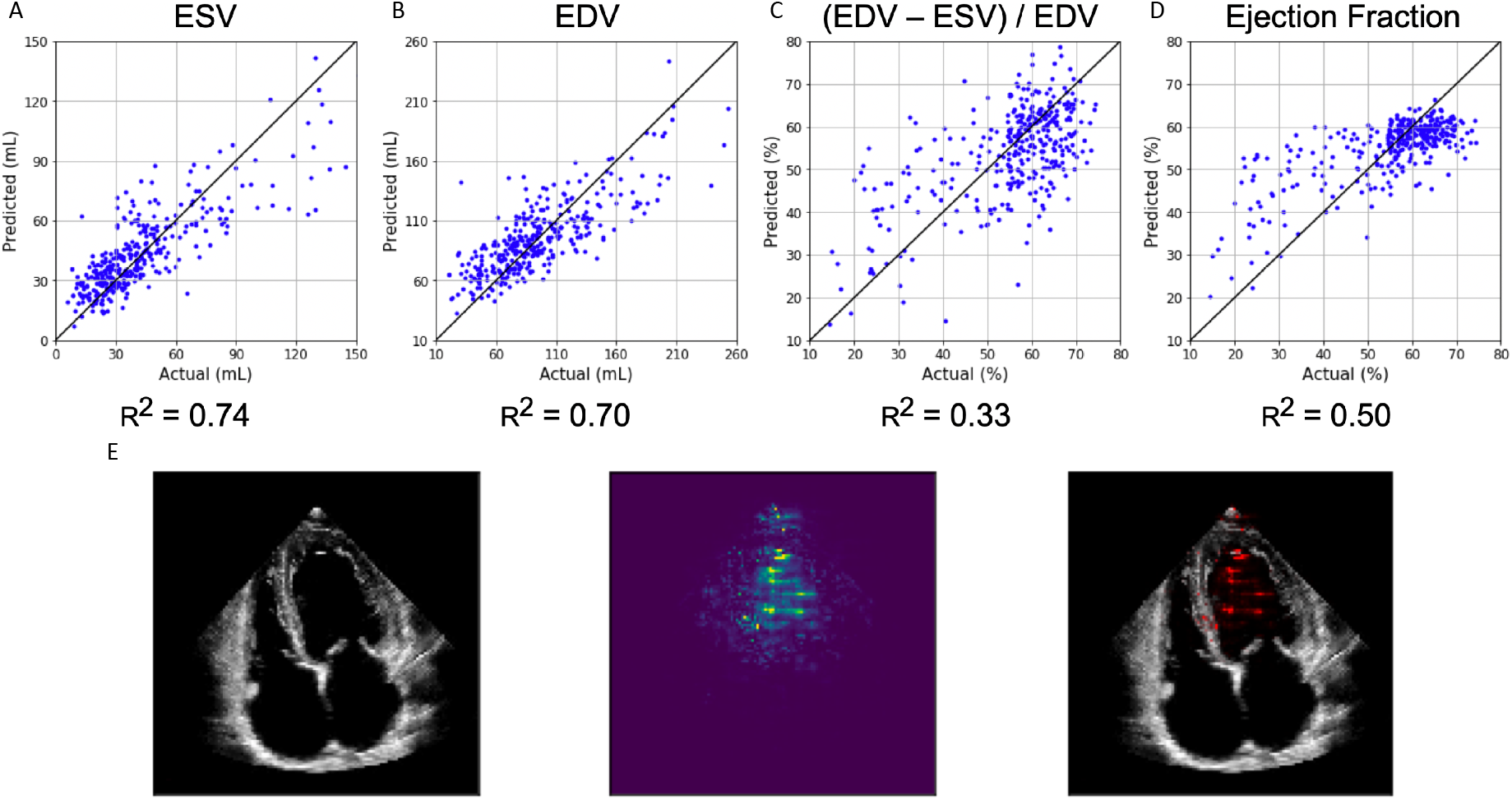
EchoNet performance for A) predicted left ventricular end systolic volume, B) predicted end diastolic volume, C) calculated ejection fraction from predicted ESV and EDV, and D) predicted ejection fraction. E) Input image, interpretation, and overlap for ejection fraction model.

### Predicting systemic cardiovascular risk factors

With good performance in identifying local structures and estimating volumetric measurements of the heart, we sought to determine if EchoNet can also identify systemic phenotypes that modify cardiovascular risk. Previous work has shown that deep convolutional neural networks have powerful capacity to aggregate the information on visual correlations between medical imaging data and systemic phenotypes.^12^ EchoNet predicted systemic phenotypes of age (*R*^2^ = 0.46, MAE = 9.8 y, mean prediction MAE = 13.4 y), sex (AUC = 0.88), weight (*R*^2^ = 0.56, MAE = 10.7 Kg, mean prediction MAE = 15.4 Kg), and height (*R*^2^ = 0.33, MAE = 0.07 m, mean prediction MAE = 0.09 m) with similar performance to previous predictions of cardiac specific features (Fig. 4A). It is recognized that characteristics such as heart chamber size and geometry vary by age, sex, weight, and height^27, 28^, however human interpreters cannot predict these systemic phenotypes from echocardiogram images alone.

**Figure 4.**
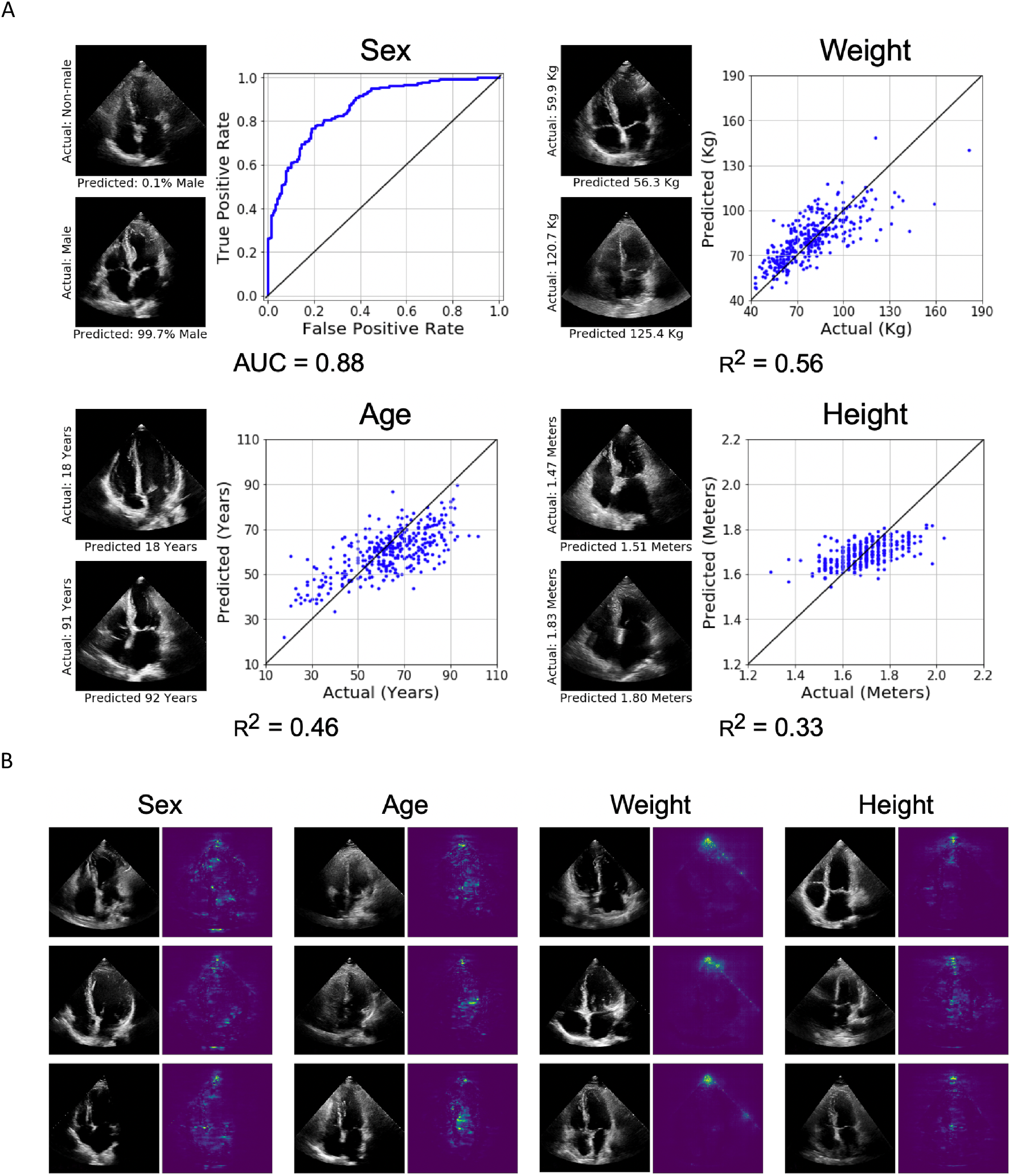
A. EchoNet performance for prediction of four systemic phenotypes (sex, weight, height and age) using apical-4-chamber view images. B. Interpretation of systemic phenotype models with representative positive examples shown side-by-side with regions of interest.

Lastly, we used the same gradient-based sensitivity map methods to identify regions of interest for models predicting systemic phenotypes difficult for human experts to predict. These regions of interest for these models tend to be more diffuse, highlighting the models for systemic phenotypes do not rely as much on individual features or local regions (Fig. 4B). The interpretations for models predicting weight and height had particular attention on the apex of the scanning sector, suggesting information related to the thickness and characteristics of the chest wall and extra-cardiac tissue was predictive of weight and height.

## Discussion

In this study, we show that deep convolutional neural networks trained on standard echocardiograms images can identify local features, human interpretable metrics of cardiac function, and systemic phenotypes such as patient age, sex, weight, and height. Our models achieved high prediction accuracy for tasks readily performed by human interpreters, such as estimating ejection fraction and chamber volumes and identifying of pacemaker leads, as well as for tasks that would be challenging for human interpreters, such as predicting systemic phenotypes from images of the heart alone. In addition to showing the predictive performance of our methods, we validate the model’s predictions by highlighting important biologically-plausible regions of interest that correspond to each interpretation. These results represent a step towards automated image evaluation of echocardiograms through deep learning. We believe this research could supplement future approaches to screen for subclinical cardiovascular disease and develop systems for personalized risk stratification. In addition, we believe the approach employed in our work, using interpretability frameworks to identify regions of interest for challenging, human-unexplainable phenotypes, may lay additional groundwork toward understanding human physiology and deep learning medical applications in medical imaging.

Previous studies have shown that medical imaging can predict cardiovascular risk factors including age, gender and blood pressure as even non-cardiac imaging displays organ-system specific manifestations of systemic phenotypes.^12^ Our results identify another avenue of detecting systemic phenotypes through organ-system specific imaging. These results are supported by previous studies that showed population level normative values for the chamber sizes of cardiac structures as participants vary by age, sex, height, and weight.^27, 28^ Age related changes in the heart, in particular changing chamber sizes and diastolic filling parameters, have been well characterized^29, 30^, and our study builds upon this body of work to demonstrate that these signals are present to allow for prediction of these phenotypes to a degree of precision not previously reported.

In addition to chamber size, extracardiac characteristics as well as additional unlabeled features, are incorporated in our models to predict patient systemic phenotypes. The area closest to the transducer, representing subcutaneous tissue, chest wall, lung parenchyma and other extracardiac structures are highlighted in the weight and height prediction models. These interpretation maps are consistent with prior knowledge that obese patients often have challenging image acquisition^31, 32^, however it is surprising the degree of precision it brings to predicting height and weight. Prediction of these systemic phenotypes suggest that imaging based predictions of mortality and life expectancy could have high predictive value as systemic phenotypes of age, sex, and body mass index are highly correlated with cardiovascular outcomes and overall life expectancy^33–35^.

Retrospective review of predictions by our model suggest human-interpretable features that show biologic plausibility. Clinical review of images predicted to be of younger patients show preference for small atria and is consistent with prior studies showing age-related changes to the left atrium^30, 36^. The feedback loop between physician and machine learning model with clinician review of appropriate and inappropriately predicted images can assist in greater understanding of normal variation in human echocardiograms as well as identify features previously neglected by human interpreters. Understanding misclassifications, such as patients with young biological age but high predicted age, and further investigation of extreme individuals can potentially help identify subclinical cardiovascular disease and better understand the aging process.

Previous studies of deep learning on medical imaging focused on resource-intensive imaging modalities common in resource-rich settings^37, 38^ or sub-speciality imaging with focused indication.^12, 13, 16^ These modalities often need retrospective annotation by experts as the clinical workflow often does not require detailed measurements or localizations. Echocardiography is one of the most frequently using imaging studies in the United States^39^ and often uses structured reporting, making advances in deep learning in echocardiography particularly applicable and generalizable. Standardization and automation of echocardiography through deep learning can make cardiovascular care more readily available. With point-of-care ultrasound is being more frequently used by an increasing number of physicians, ranging from emergency room physicians, internists, to anaesthesiologists, and deep learning on cardiac ultrasound images can provide accurate predictions and diagnoses to an even wider range of patients.

In summary, we provide evidence that deep learning can reproduce common human interpretation tasks and leverage additional information to predict systemic phenotypes that could allow for better cardiovascular risk stratification. We used interpretation methods that could feedback relevant regions of interest for further investigation by cardiologists to better understand aging and prevent cardiovascular disease. Our work could enable assessment of cardiac physiology, anatomy, and risk stratification at the population level by automating common workflows in clinical echocardiography and democratize expert interpretation to general patient populations.

## Methods

### Dataset

The Stanford Echocardiography Database contains images, physician reports, and clinical data from patients at Stanford Hospital who underwent echocardiography in the course of routine care. The accredited echocardiography laboratory provides cardiac imaging to a range of patients with a variety of cardiac conditions including atrial fibrillation, coronary artery disease, cardiomyopathy, aortic stenosis, and amyloidosis. For this study, we used 3312 comprehensive non-stress echocardiography studies obtained between January 2017 and December 2018, and split the patients into training and validation cohorts. Videos of standard cardiac views, color Doppler videos, and still images comprise each study and is stored in Digital Imaging and Communications in Medicine (DICOM) format. The videos were sampled at a rate of 10 frames per second to obtain 1,624,780 unscaled 299×299 pixel images. For each image, information pertained to image acquisition, identifying information, and other information outside the imaging sector was removed through masking. Human interpretations from the physician-interpreted report and clinical features from the electronic medical record were matched to each echocardiography study for model training. This study was approved by the Stanford University IRB.

### Model

We chose a convolutional neural network (CNN) architecture that balances network width and depth in order to manage the computational cost of training. We used the architecture based on Inception-Resnet-v1^10^ to predict all of our phenotypes. This architecture has strong performance on benchmark datasets like ILSVR2012 image recognition challenge (Imagenet)^40^ and is computationally efficient compared to other networks^41^. Using architectures pretrained on ImageNet data set did not result in any performance improvement.

For each prediction task, one CNN architecture was trained on individual frames from each echocardiogram video with output labels that were extracted either from the electronic medical record or from the physician report. From each video, we sampled 20 frames (one frame per 100 milliseconds) starting from the first frame of the video. The final prediction was performed by averaging all the predictions from individual frames. Several alternative methods were explored in order to aggregate frame-level predictions into one patient-level prediction and did not yield better results compared to simple averaging. We also investigated multi-task learning—sharing some of the model parameters while predicting across the different phenotypes—and this did not improve the model performance.

Model training was performed using the TensorFlow library^42^ which is capable of utilizing parallel-processing capabilites of Graphical Processing Units (GPUs) for fast training of deep learning models. We chose Adam optimizer as our optimization algorithm which is computationally efficient, has little memory usage, and has shown superior performance in many deep learning tasks^43^. As our prediction loss, we used cross-entropy loss for classification tasks and squared error loss for regressions tasks along with using weight-decay regularization loss to prevent over-fitting^44^. We investigated other variants of prediction loss (absolute loss, Huber loss^45^ for regression and Focal loss^46^ for classification), and they did not improve performance. For each prediction task, we chose the best performing hyper-parameters using grid search (24 models trained for each task) to optimize learning rate and weight decay regularization factor. In order to perform model selection, for each tasks, we split the training data into training and validation set by using 10% of train data as a held-out validation set in; the model with the best performance on the validation set is then examined on the test set to report the final performance results. After the models were trained, they were evaluated on a separate set of test frames gathered from echocardiogram studies of 337 other patients (separate from training data patients) with similar demographics.

### Data augmentation

Model performance improved with increasing input data sample size. Our experiments suggested additional relative improvement with increase in the number of patients represented in the training cohort compared to oversampling of frames per patient. Data augmentation using previously validated methods^47, 48^, also greatly improving generalization of model predictions by reducing over-fitting on the training set. Through the training process, at each optimization step each training image is transformed through geometric transformations (such as flipping, reflection, and translation) and changes in contrast and saturation. As a result, the training data set is augmented into a larger effective data set. In this work, mimicking variation in echocardiography image acquisition, we used random rotation and random saturation augmentation for data augmentation (Fig. **??**C). During each step of stochastic gradient descent in the training process, we randomly sample 24 training frames, and we perturb each training frame with a random rotation between −20 to 20 degrees and with adding a number sampled uniformly between −0.1 to 0.1 to image pixels (pixels values are normalized) to increase or decrease brightness of the image. Data augmentation results in improvement for all of the tasks; between 1 − 4% improvement in AUC metric for classification tasks and 2 − 10% improvement in *R*^2^ score for regression tasks.

### Cardiac view selection

We first tried using all echocardiogram images for prediction tasks but given the size of echocardiogram studies, initial efforts struggled with long training times, poor model convergence, and difficulty with model saturation. With the knowledge that, in a single comprehensive echocardiography study, the same cardiac structures are often visualized from multiple views to confirm and corroborate assessments from other views, we experimented with model training using subsets of images by cardiac view. As described in Fig. **??**B, a selection of the most common standard echocardiogram views were evaluated for model performance. Images from each study were classified using a previously described supervised training method^5^. We sought to identify the most information-rich views by training separate models on the subsets of dataset images of only one cardiac view. Training a model using only one cardiac view results in one order of magnitude reduction of training time and computational cost with the benefit of maintaining similar predictive performance when information-rich views were used. Given the favorable balance of performance to computational cost as well as prior knowledge on which views most cardiologists frequently prioritize, we chose the apical-4-chamber view as the input training set for subsequent experiments on training local features, volumetric estimates and systemic phenotypes.

### Interpretability

Interpretability methods for deep learning models have been developed to explain the predictions of the black-box deep neural network. One family of interpretations methods are the sensitivity map methods that seek to explain a trained model’s prediction on a given input by assigning a scalar importance score to each of the input features or pixels. If the model’s input is an image, the resulting sensitivity map could be depicted as a two-dimensional heat-map with the same size as the image where more important pixels of the image are brighter than other pixels. The sensitivity map methods compute the importance of each input feature as the effect of its perturbation on model’s prediction. If the pixel is not important, the change should be small and vice versa.

Introduced by Baehrens et al.^49^ and applied to deep neural networks by Simonyan et al. ^50^, the simplest way to compute such score is to have a first-order linear approximation of the model by taking the gradient of the output with respect to the input; the weights of the resulting linear model are the sensitivity of the output to perturbation of their corresponding features (pixels). More formally, given the *d*-dimensional input **x**_*t*_ ∈ ℝ^*d*^ and the model’s prediction function *f* (.), the importance score of the *j*’th feature is |∇_**x**_*f*(**x**_*t*_)_*j*_|. Further extensions to this gradient method were introduced to achieve better interpretations of the model and to output sensitivity maps that are perceptually easier to understand by human users: LRP^51^, DeepLIFT^52^,Integrated Gradients^53^, and so forth. These sensitivity map methods, however, suffer from visual noise^25^ and sensitivity to input perturbations.^54^. SmoothGrad^25^ method alleviates both problems^55^ by adding white noise to the image and then take the average of the resulting sensitivity maps. In this work, we use SmoothGrad with the simple gradient method due to its computational efficiency. Other interpretation methods including Integrated Gradients were tested but did not result in better visualizations.

### Lessons from model training and experiments

EchoNet performance greatly improved with efforts to augment data size, homogenize input data, and with optimize model training with hyperparameter search. Our experience shows that increasing number of unique patients in the training set can significantly improve the model, more so than increasing the sampling rate of frames from the same patients. Homogenizing the input images by selection of cardiac view prior to model training greatly improved training speed and decreased computational time without significant loss in model performance. Finally, we found that results can be significantly improved with careful hyperparameter choice; between 7 − 9% in AUC metric for classification tasks and 3 − 10% in *R*^2^ score for regression tasks.

## Data Availablity

The data comes from medical records and imaging from Stanford Healthcare and is not publicly available. The de-identified data is available from the authors upon reasonable request and with permission of the institutional review board.

## Acknowledgements

This work is supported by the Stanford Translational Research and Applied Medicine pilot grant and an Stanford Artificial Intelligence in Imaging and Medicine Center seed grant. D.O. is supported by the American College of Cardiology Foundation / Merck Research Fellowship. A.G. is supported by the Stanford-Robert Bosch Graduate Fellowship in Science and Engineering. J.Z. is supported by a Chan-Zuckerberg Biohub Fellowship.

